# The path to re-evolve cooperation is constrained in *Pseudomonas aeruginosa*

**DOI:** 10.1101/163048

**Authors:** Elisa T. Granato, Rolf Kümmerli

**Affiliations:** Department of Plant and Microbial Biology, University of Zurich, Zurich, Switzerland

**Keywords:** cooperation, siderophores, cheating, iron, Pseudomonas, *pvdS*

## Abstract

**Background:** A common form of cooperation in bacteria is based on the secretion of beneficial metabolites, shareable as public good among cells at the group level. Because cooperation can be exploited by “cheat” mutants, which contribute less or nothing to the public good, there has been great interest in understanding the conditions required for cooperation to remain evolutionarily stable. In contrast, much less is known about whether cheats, once fixed in the population, are able to revert back to cooperation when conditions change. Here, we tackle this question by subjecting experimentally evolved cheats of *Pseudomonas aeruginosa*, partly deficient for the production of the iron-scavenging public good pyoverdine, to conditions previously shown to favor cooperation.

**Results:** Following approximately 200 generations of experimental evolution, we screened 720 evolved clones for changes in their pyoverdine production levels. We found no evidence for the re-evolution of full cooperation, even in environments with increased spatial structure, and reduced costs of cooperation – two conditions that have previously been shown to maintain cooperation. In contrast, we observed selection for complete abolishment of pyoverdine production. The patterns of complete trait degradation were likely driven by “cheating on cheats” in unstructured, iron-limited environments where pyoverdine is important for growth, and selection against a maladaptive trait in iron-rich environments where pyoverdine is superfluous.

**Conclusions:** Our study shows that the path to re-evolve cooperation seems constrained. One reason might be that the number of mutational targets potentially leading to reversion is limited. Alternatively, it could be that the selective conditions required for revertants to spread from rare are much more stringent than those needed to maintain cooperation.

## BACKGROUND

Bacterial life predominantly takes place in diverse communities, where individual cells are constantly surrounded by neighbors. While high cell density and diversity can create strong competition in the struggle for nutrients and space [1,2], it can also promote stable networks of cooperation [3,4]. A common way for bacteria to cooperate is through the secretion of nutrient-scavenging metabolites, which are shared as “public goods” in the community. Public goods cooperation is thought to increase nutrient uptake rate, and results in the costs and benefits of public goods being shared among producer cells. Although beneficial for the collective as a whole, public goods cooperation can select for “social cheats”: mutants that lower or abolish their investment into public good production, but still reap the benefits of nutrient uptake [5,6].

The undermining of public goods cooperation by cheats has spurred an entire field of research, examining the conditions required for cooperation to be maintained in the population. In contrast, the question of how public goods cooperation evolves in the first place has received much less attention. The main question is: will the conditions that have been shown to maintain cooperation also promote the evolution of cooperation? Here, we tackle this question by examining whether bacteria that have evolved low levels of cooperation in a previous experiment can evolve back to normal levels of cooperation under conditions that are known to be favorable for cooperation. We use pyoverdine, an iron-scavenging siderophore secreted by the opportunistic pathogen *Pseudomonas aeruginosa*, as our model cooperative trait. Pyoverdine is the main siderophore of *P. aeruginosa*, and is secreted into the environment in response to iron limitation. Pyoverdine acts as a shareable public good that can be exploited by non-producing cheats that possess the matching receptor for uptake [7,8].

We consider three factors that could determine whether cooperation can re-evolve or not. The first factor is the spatial structure of the environment. Previous work revealed that increased spatial structure maintains cooperation because it reduces pyoverdine diffusion and cell dispersal. In other words, spatial structure ensures that pyoverdine sharing occurs predominantly among producer cells [9,10]. The second factor involves the relative costs and benefits of pyoverdine production [8,11]. In the absence of significant spatial structure, it was shown that cheats enjoyed highest relative fitness advantages under severe iron limitation when pyoverdine is expressed at high levels (i.e. high costs). Conversely, cooperation was maintained at intermediate iron limitation when pyoverdine is still important for growth, yet its investment is reduced (i.e. lower costs). Finally, we examine whether the genetic background of cheats is an important determinant of whether cooperation can re-evolve. Previous studies [7,12; Granato ET, Ziegenhain C & Kümmerli R, unpublished] observed the evolution of two types of cheats with greatly decreased pyoverdine production. The first type of cheat has a point mutation in *pvdS*, the gene encoding the sigma factor regulating pyoverdine production [13], whereas the second type of cheat has a point mutation in the promoter region of *pvdS*. While the two types of mutations might differ in their likelihood to revert back to cooperation, both could principally do so, because their pyoverdine biosynthesis cluster is intact [14], and a single point mutation in regulatory elements could lead to reversion.

We conducted experimental evolution in replicated populations with the two types of pyoverdine deficient strains across three levels of iron limitations and two habitats, differing in their level of spatial structuring. Based on social evolution theory, we predict the reversion to full cooperation whenever Hamilton’s rule [15] – *rB > C* – is satisfied. While *r* is the relatedness between the actor and the recipient, *C* is the cost to the actor performing cooperation, and *B* is the benefit gained by the individual receiving cooperation. In our treatments, we vary *r* by manipulating the degree of spatial structure and *C*/*B* by manipulating the level of iron limitation. Accordingly, we predict that increased spatial structure and/or moderate investments into pyoverdine production should be most conducive for the re-evolution of cooperation. Moreover, we also envisage the possibility of pyoverdine production to degrade even further. This seems plausible because the mutated clones still produce some pyoverdine, and thus, there is room for further exploitation by *de novo* mutants that make even less. We predict this to happen under low spatial structure, and high pyoverdine investment levels. Finally, pyoverdine could also be degraded due to disuse [16], especially under conditions of high iron availability where pyoverdine is not required.

## RESULTS

### Characterization of the ancestral pyoverdine deficient strains

We first characterized the strains *pvdS*_gene and *pvdS*_prom for their pyoverdine production and growth dynamics (Fig. 1) before they were subjected to experimental evolution (Fig. 2). These two mutants themselves spontaneously arose and spread during a previous experimental evolution study (Granato ET, Ziegenhain C & Kümmerli R, unpublished). Their entire genomes had been re-sequenced and analyzed. Those analyses revealed that both *pvdS*_gene and *pvdS*_prom carried non-synonymous mutations that are directly associated with their reduced pyoverdine investment levels (Fig. 1a). Strain *pvdS*_gene has a point mutation (G>C) in the *pvdS* gene that leads to an amino acid change (Met135Ile), and thus to a modified iron-starvation sigma factor PvdS. A modified PvdS presumably has lower affinity to the RNA-polymerase, a complex that directly controls the expression of the non-ribosomal peptide synthesis machinery required to build pyoverdine. Strain *pvdS*_prom carries a point mutation (G>T) in the consensus sequence of the −35 element in the promoter region upstream of *pvdS*. This mutant produces a wildtype sigma factor, but the transcription rate of PvdS is likely reduced.

**Fig. 1:**
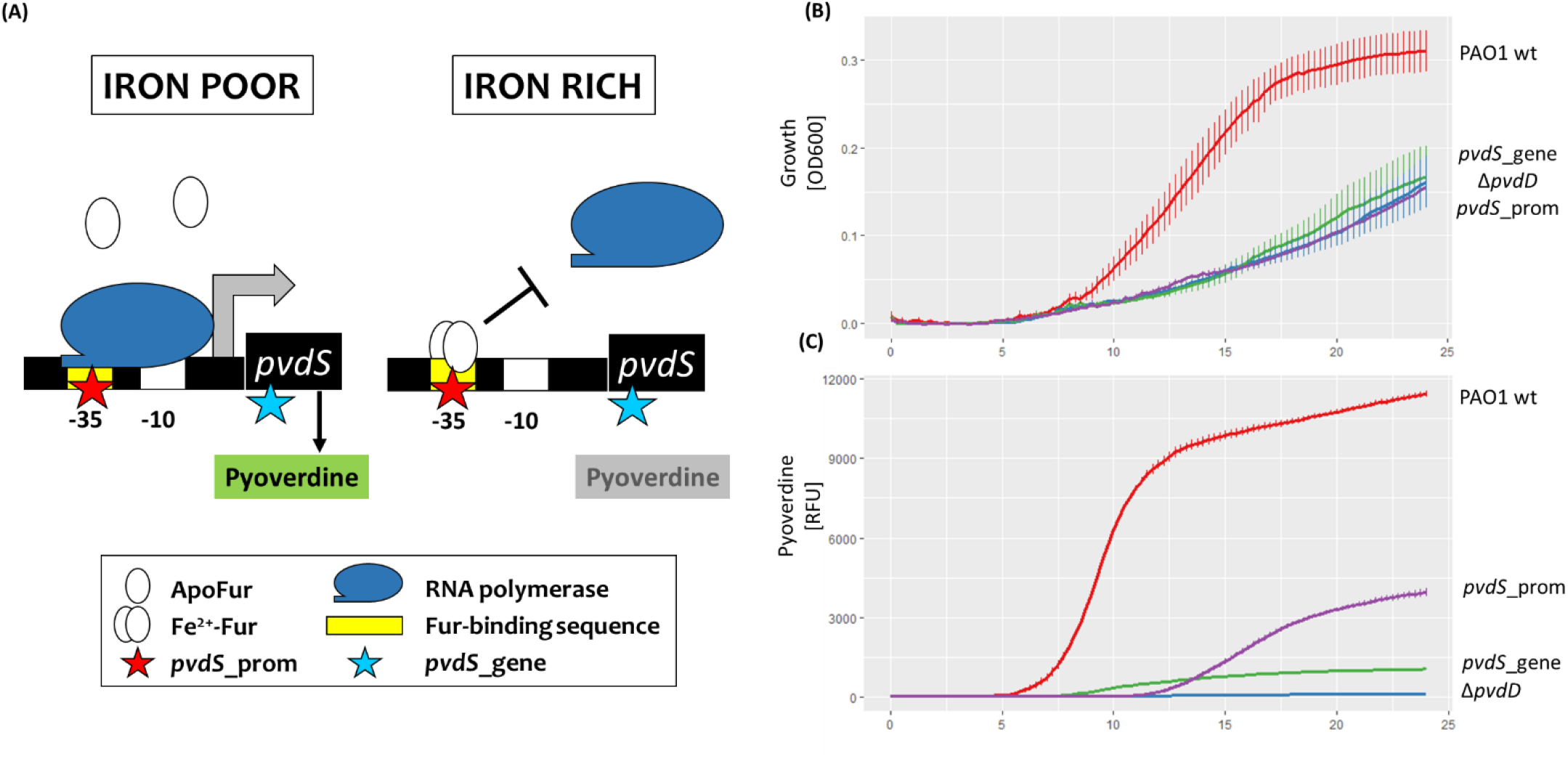
Evolved clones of *P. aeruginosa* show impaired growth and pyoverdine production. Schematic representation of *pvdS* regulation under iron-poor and iron-rich conditions. When iron is limited, *pvdS* is transcribed and upregulates pyoverdine biosynthesis. When iron levels in the cytoplasm are sufficient, Fur (ferric-<u>u</u>ptake <u>r</u>egulator) builds a complex with Fe^2+^, which then binds to the *pvdS*-promoter site and inhibits transcription. Stars indicate SNPs in the mutant strains *pvdS*_prom (red) and *pvdS*_gene (blue). **(B+C)** A *P. aeruginosa* wildtype strain (PAO1 wt) and three different mutants with deficient pyoverdine production were grown in iron-limited media at 37 °C for 24 hours. Y axis shows **(B)** optical density measured at 600 nm or **(C)** pyoverdine-specific fluorescence (emission|excitation 400 nm|460 nm). X axis shows time in hours. *ΔpvdD*: engineered knock-out mutant carrying an in-frame deletion of *pvdD*, encoding a part of the pyoverdine synthesis pathway. *pvdS*_gene: evolved mutant with single point mutation in *pvdS*, encoding the iron-starvation sigma factor PvdS. *pvdS*_prom: evolved mutant with a single point mutation in the promoter region of *pvdS*. Graph depicts means and standard errors based on four independent replicates per strain.

**Fig. 2:**
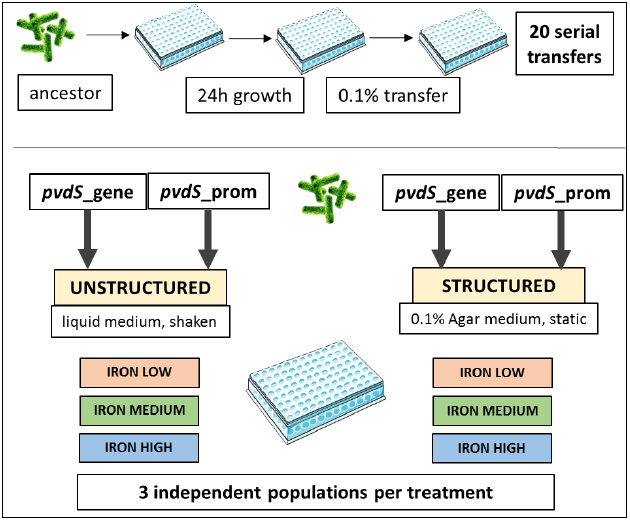
Experimental evolution setup. Two mutant strains deficient in pyoverdine production, *pvdS*_gene and *pvdS*_prom, were allowed to evolve independently from each other and under different conditions. *pvdS*_gene carries a single point mutation in *pvdS*, encoding the iron-starvation sigma factor PvdS, while *pvdS*_prom carries a single point mutation in the promoter region of *pvdS*. The six environments used for experimental evolution differed both in their level of spatial structure (unstructured| structured) and in their iron content (“iron low”: iron chelator only; “iron medium”: iron chelator + 1 μM FeCl_3_; “iron high”: iron chelator + 40 μM FeCl_3_). Each ancestral strain was serially transferred in each of the six media in threefold replication, resulting in a total number of 36 independently evolved populations. Image sources: Servier Medical Art (multiwell plate); depositphotos.com (bacteria).

Both of these mutations show strong defects in pyoverdine production and growth under iron-limited conditions (Fig. 1b+c). Pyoverdine production of the *pvdS*_gene strain was only 9.4 ±0.1 % (mean ± SE) compared to the wildtype strain PAO1 (measured after 24 hours), and characterized by a low but steady production rate (Fig. 1c). While pyoverdine production was also reduced in *pvdS*_prom (34.7 ± 1.4 % relative to the ancestral wildtype strain), the production dynamic differed from *pvdS*_gene. The *pvdS*_prom strain had an extended phase, where no pyoverdine is produced, followed by a phase with a considerable production rate (Fig. 1c). Both mutant strains displayed substantial growth impairments, comparable to that of a constructed pyoverdine knockout (Fig. 1b). This indicates that the production of higher amounts of pyoverdine would be advantageous.

### Further degradation and not re-evolution of pyoverdine production prevails

Following 20 days (approx. 200 generations) of experimental evolution in six different environments (2 different spatial structures x 3 different iron concentrations; Fig. 2), we screened 720 clones for their evolved levels of pyoverdine production and growth under iron limitation (Fig. 3). For each clone, we then calculated the per capita pyoverdine production (pyoverdine fluorescence divided by OD_600_). Under the conditions of this assay, the ancestral strains *pvdS*_gene and *pvdS*_prom displayed 17.4 and 28.5 % (unstructured|structured), and 59.9 and 83.2 % (unstructured|structured) of the wildtype PAO1 pyoverdine production levels, respectively. Among the evolved clones, there were only very few (n = 5; 0.69%) that exhibited considerably increased pyoverdine production levels (Fig. 3), indicating that reversion to higher levels of cooperation is rare. In contrast, we found a considerable number of clones (n = 29; 4.03 %) that showed either a complete abolishment or a further substantial reduction in pyoverdine production during evolution.

**Fig. 3:**
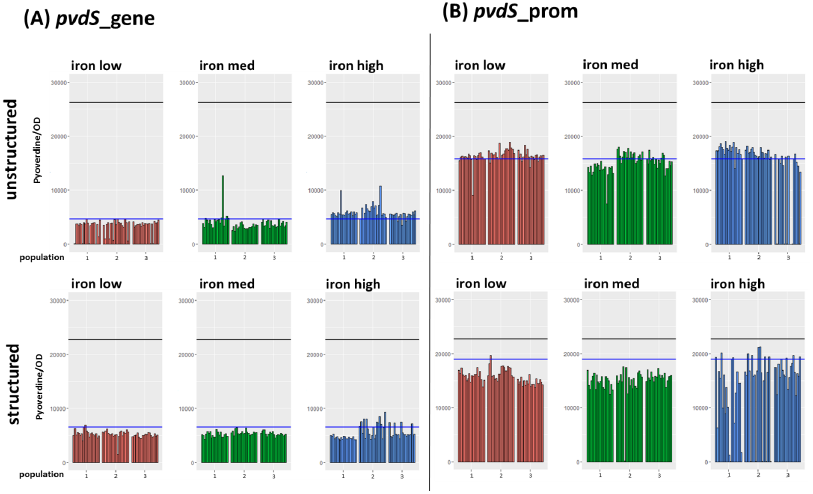
Changes in pyoverdine production after experimental evolution in different environments. Two mutants with abnormal pyoverdine production were allowed to evolve in different media, and pyoverdine production under iron-limitation was subsequently measured in 1200 evolved clones. Environments differed in their level of spatial structure (structured|unstructured) and in their iron content (“iron low”: iron chelator only; “iron med”: iron chelator + 1 μM FeCl_3_; “iron high”: iron chelator + 40 μM FeCl_3_). **(A)** Clones evolved from the low-producer *pvdS*_gene, a mutant with a single point mutation in *pvdS*, encoding the iron-starvation sigma factor PvdS. **(B)** Clones evolved from the low-producer *pvdS*_prom, a mutant with a single point mutation in the promoter region of *pvdS*. Y axes show pyoverdine-specific fluorescence divided by growth (optical density at 600 nm) after 24 h of incubation. X axes show independent replicate populations the clones evolved in. Each bar represents a single measurement per evolved clone. The black line denotes the average wildtype production level in the same assay, while the blue line denotes the average production level of the respective low-producing ancestor.

There was an interaction between the genetic background and the environmental conditions under which these non-and extremely low pyoverdine-producing mutants appeared. In the *pvdS*_gene background, they appeared exclusively under low iron conditions, and were significantly more prevalent in unstructured compared to structured environments (Fisher’s exact test, p = 0.012; Table 1). Since pyoverdine is important for growth under these iron-limited conditions, yet can be exploited in unstructured environments, this pattern suggests that the non-and extremely low pyoverdine-producing clones are cheats, which spread because they exploited the little amount of pyoverdine produced by *pvdS*_gene. In the *pvdS*_prom background, meanwhile, non-and extremely low-producers appeared almost exclusively under high iron conditions (Fisher’s exact test, p < 0.001), but independently of the spatial structure (Fisher’s exact test, p = 0.78; Table 1). This pattern indicates that pyoverdine production was eroded due to disuse in iron-rich environments.

**Table 1.** Frequency of non-and low-producing strains per treatment

| ancestor | *pvdS_gene* |  |  |  |  |  | *pvdS_prom* |  |  |  |  |  |
| --- | --- | --- | --- | --- | --- | --- | --- | --- | --- | --- | --- | --- |
| environment | structured |  |  | unstructured |  |  | structured |  |  | unstructured |  |  |
| iron<sup>a</sup> | low | med | high | low | med | high | low | med | high | low | med | high |
| NLPs<sup>b</sup> | 1 | 0 | 0 | 11 | 0 | 0 | 0 | 1 | 9 | 0 | 0 | 7 |
| rest<sup>c</sup> | 59 | 60 | 60 | 49 | 60 | 60 | 60 | 59 | 51 | 60 | 60 | 53 |

<sup>a</sup> low, medium (med) or high iron availability; see methods for details
<sup>b</sup> non- or low-producers, based on initial screening; corresponds to data shown in Fig. 3
<sup>c</sup> clones not in the NLP category

### In-depth analysis of a subset of evolved clones confirms selection against pyoverdine

Since the large screen of 720 clones was based on a single replicate per clone (Fig. 3), we subjected the 34 clones with a putatively altered pyoverdine phenotype to a replicated in-depth phenotypic screen. We further included 23 clones with apparently unaltered pyoverdine phenotypes. For clones with the *pvdS*_gene background, we could confirm the phenotype of all clones that showed a further decrease in pyoverdine production (Fig. 4a). In fact, pyoverdine production virtually absent in all of them. Conversely, we could only confirm the phenotype of two of the three mutants with putatively increased pyoverdine production, and even for the confirmed ones, the observed increase was marginal (Fig. 4b). We obtained similar confirmation patterns for clones with the *pvdS*_prom background: confirmation rate was only high for clones with reduced but not for those with increased pyoverdine production levels (Fig. 4c+d). Finally, when examining the clones with a putatively unaltered pyoverdine, we found that 61 % (14 out of 23) of these clones indeed had a phenotype equal to their ancestral strain, whereas 35 % (8 out of 23) of the clones had pyoverdine production slightly but significantly reduced (Fig. S1 in Additional File 1). Taken together, these results confirm the patterns of our extensive screen (Fig. 3): there was selection to further reduce pyoverdine production, but no restoration of cooperation.

**Fig. 4:**
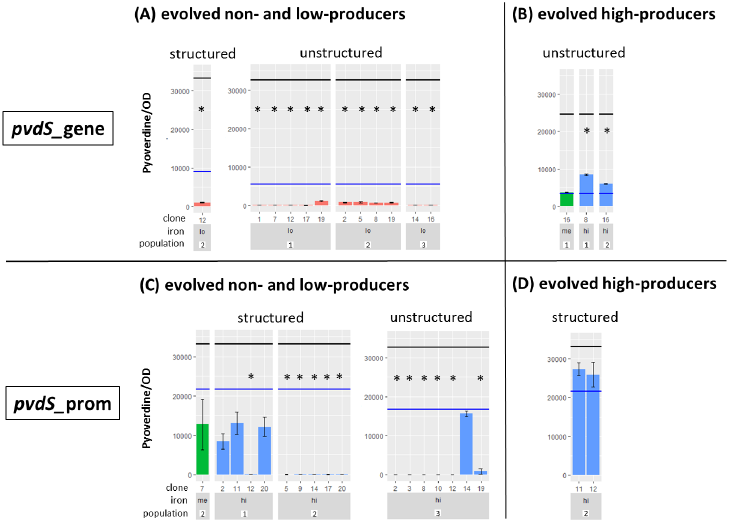
Confirmed pyoverdine phenotypes in selected clones. Evolved clones with changes in pyoverdine production were re-tested to confirm their phenotype. Pyoverdine production was measured in iron-limited media. **(A)** Clones evolved from the low-producer *pvdS*_gene with low production levels in initial screening. **(B)** Clones evolved from the low-producer *pvdS*_gene with high production levels in initial screening. **(C)** Clones evolved from the low-producer *pvdS*_prom with low production levels in initial screening. **(D)** Clones evolved from the low-producer *pvdS*_prom with high production levels in initial screening. Y axes show pyoverdine-specific fluorescence divided by growth (optical density at 600 nm) after 24 h of incubation. X axes show independent replicate populations the clones evolved in and iron availability during experimental evolution. Bars represent mean values of three replicates per evolved clone. Error bars denote standard error of the mean. The black line represents the average wildtype production level in the same assay, while the blue line denotes the average production level of the respective low-producing ancestor. Bars are coloured by iron availability during evolution: red = low iron; green = medium iron; blue = high iron. We used one-way ANOVAs with Tukey’s post-hoc test for comparisons relative to the low-producing ancestor. Asterisks indicate a significant difference (p<0.05) from the low-producing ancestor.

### Evolved pyoverdine phenotypes are not based on further mutations in *pvdS*

We anticipated that both restoration and further reduction of pyoverdine production could be caused by additional mutations in the *pvdS* gene or its promoter. However, we found no support for this hypothesis when sequencing this genetic region for the subset of 57 clones described above (Table S1 in Additional File 1). All clones had retained the original, ancestral mutation inherited from their respective low-producing ancestor (SNP in the *pvdS* gene itself for *pvdS*_gene, SNP in the *pvdS* promoter region for *pvdS*_prom). Additionally, one clone from the *pvdS*_gene line gained an additional SNP in the *pvdS* promoter region, which however did not affect its phenotype. No additional mutations were found in any of the clones, indicating that the observed changes in pyoverdine production either represent an entirely phenotypic change, or are caused by mutations in regions other than *pvdS*.

## DISCUSSION

Numerous studies used microbial systems to address a key question in evolutionary biology: how can cooperation be maintained in the face of cheats that exploit the cooperative acts performed by others [17–19]. Conversely, the question of what happens after a cheat has become fixed in the population has received much less attention. Would it be possible that cooperation re-evolves if environmental conditions and thus selection pressures change [20,21]? To tackle this question, we performed experimental evolution with *P. aeruginosa* cheat strains (mutants that produced greatly reduced amounts of the iron-scavenging public good pyoverdine), which had the potential to revert back to a full cooperative phenotype by a single point mutation. Despite this favourable genetic predisposition, we never observed reversion to cooperation, even under conditions that had previously been identified as being favourable for cooperation. Instead, we observed the emergence of mutants that completely abolished pyoverdine production, and their frequency of appearance depended on both their genetic background and the environmental conditions. Taken together, our study highlights that the re-evolution of cooperation might be constrained in bacteria.

We can think of at least two reasons why there was no reversion from cheats back to cooperators. At the mechanistic level, it might be that the likelihood of acquiring a mutation that leads to reversion was simply too low. It is well known that mutations causing a loss of function are disproportionately more likely to occur than mutations resulting in a gain of function [22]. In the context of our experiment, re-evolution of pyoverdine production could have happened by a reversion to the ancestral PAO1 genotype (i.e. reversing the point mutation in the *pvdS* region) or by a compensatory mutation in *pvdS* or another regulatory element. Clearly, the number of mutational targets that would lead to reversion seem limited, and thus mutation supply might have been too low for revertants to arise.

At the ultimate level, it might be that we have not chosen the appropriate environmental conditions that would select for reversion. According to Hamilton’s rule, we would expect selection for reverted cooperators when relatedness is relatively high and/or when the cost-to-benefit ratio of cooperation is relatively low. Although we have implemented experimental conditions promoting significant relatedness (through limited cell mixing in spatially structured environments) and reduced costs of pyoverdine production (at intermediate iron limitation), the chosen conditions were apparently not favourable enough to select for the re-evolution of cooperation. At first glance, this seems surprising because the chosen conditions have previously been shown to prevent the spreading of cheats and to maintain cooperation [8,10,23]. Our findings thus suggest that the conditions for the evolution of cooperation are more stringent than those for the maintenance of cooperation. Indeed, social evolution theory predicts cooperation to be maintained when *rb* = *c* (rare cheats cannot invade), while Hamilton’s rule *rb* > *c* must be met for cooperation to evolve. The fulfilment of this latter condition might require specific conditions (e.g. very high relatedness), as reverted cooperators would have to invade from extreme rarity, while being surrounded by clones exploiting any pyoverdine molecule diffusing away from the producer.

Instead of reversion to cooperation, we observed selection for mutants that further reduced or completely abolished pyoverdine production (Fig. 3+4). Intriguingly, the environments that promoted the spread of these mutants differed between *pvdS*_gene and *pvdS*_prom, indicating that different selection pressures can promote the same phenotype. For the *pvdS*_gene background, we found that the further degradation of pyoverdine production predominantly occurred with low spatial structure and under stringent iron limitation. As pyoverdine is important for growth under these conditions but widely shared due to mixing,we assume that these mutants spread because they cheated on the residual pyoverdine produced by the ancestral *pvdS*_gene. This finding confirms the notion that “cheating” is context-dependent, and shows that a strain that evolved as a cheat is still susceptible to further exploitation, despite its greatly reduced investment into a cooperative trait [24]. In contrast to this pattern, we observed the further degradation of pyoverdine production in the *pvdS*_prom background almost exclusively in iron-rich environments regardless of spatial structure. Because pyoverdine is not needed under iron-rich conditions, yet still expressed in low amounts [11,16], we assume that selection against pyoverdine production represents the erosion of an unnecessary trait.

We can only speculate about why the genetic background seems to matter for whether pyoverdine degradation is presumably driven by cheating or disuse. One possible explanation might reside in the different pyoverdine production profiles shown by the two strains. While *pvdS*_gene has a low but steady production rate, *pvdS*_prom delays pyoverdine production, but then produces pyoverdine at a higher rate compared to *pvdS*_gene. It could be that delaying the onset of pyoverdine production is a successful strategy to prevent the invasion of mutants with completely abolished pyoverdine production. With regard to trait erosion, it seems possible that *pvdS*_prom produces higher amounts of pyoverdine compared to *pvdS*_gene under iron-rich conditions; this would make this strain more susceptible for trait erosion because pyoverdine production is maladaptive under these conditions. Further studies are clearly needed to elucidate these pattern at both the proximate and ultimate level. The proximate level is of special interest here because the complete loss of pyoverdine production did not involve mutations in *pvdS*, which has been identified as the main target of selection for the initial reduction in pyoverdine production [7,12; Granato ET, Ziegenhain C & Kümmerli R, unpublished].

## CONCLUSIONS

Our findings indicate that the evolution of cooperation through mutational reversion seems to be constrained. Reasons for this could be linked to the low number of mutational targets available that can lead to reversion, or the stringent selective conditions required to promote the spread of revertants. Clearly, the conditions that have previously been shown to maintain cooperation are not sufficient to promote the invasion of *de novo* re-evolved cooperators from rare. While we focussed on the re-evolution of cooperation via mutations, another alternative scenario under natural conditions is that cheats may revert to cooperators through horizontal gene transfer [25,26]. This scenario has especially been advocated for cooperative traits located on plasmids [27,28]. While this is a plausible scenario for some social traits, it is unlikely to apply to siderophores, which are typically encoded on the chromosome. The insights gained from our study contribute to our understanding of the conditions necessary for a cooperative trait to evolve in microorganisms.

## METHODS

### Strains and growth conditions

We used *Pseudomonas aeruginosa* wildtype strain PAO1 (ATCC 15692) and a pyoverdine-negative mutant, both constitutively expressing GFP (PAO1-*gfp,* PAO1-Δ*pvdD*-*gfp*), as positive and negative controls for pyoverdine production, respectively. We further used PAO1-*pvdS*_gene and PAO1-*pvdS*_prom, two mutants with strongly reduced pyoverdine production, that evolved *de novo* from PAO1-*gfp* during experimental evolution in iron-limited media (2.5 gL^-1^ BactoPeptone, 3 gL^-1^ NaCl, 5 mgL^-1^ Cholesterol, 25 mM MES buffer pH = 6.0, 1mM MgSO4, 1mM CaCl2, 200 μM 2,2’-Bipyridyl (Granato ET, Ziegenhain C & Kümmerli R, unpublished)). PAO1-*pvdS*_gene carries a non-synonymous point mutation (G>C) in the *pvdS* gene that leads to an amino acid change (Met135Ile). PAO1-*pvdS*_prom carries a point mutation (G>T) in the consensus sequence of the −35 element in the promoter region upstream of *pvdS*. Both mutants constitutively express GFP. Throughout this publication, the two mutants are referred to as “*pvdS*_gene” and “*pvdS*_prom”.

For overnight pre-culturing, we used Luria Bertani (LB) medium, and incubated the bacteria under shaking conditions (190-200 rpm) for 16-18 hours. Optical density (OD) of pre-cultures was determined at a wavelength of 600 nm in a spectrophotometer. We induced strongly iron-limiting growth conditions by using casamino acids (CAA) medium (5 gL^-1^ casamino acids;1.18 gL^-1^ K_2_HPO_4_·3H_2_O; 0.25 gL^-1^ MgSO_4_·7H_2_O) supplemented with 25 mM HEPES and 400 μM of the iron chelator 2,2’-Bipyridyl. For conditions with medium or high iron availability, we further added FeCl_3_ at final concentrations of 1 μM or 40 μM, respectively. We manipulated the spatial structure of the environment by growing bacteria either in liquid medium under shaking conditions (180 rpm; unstructured environment) or in viscous medium containing 0.1% agar under static conditions (structured environment). All experiments in this study were conducted at 37°C. All chemicals were purchased from Sigma-Aldrich, Switzerland.

### Ancestral growth and pyoverdine kinetics

To measure growth and pyoverdine production kinetics of all strains in iron-limited media prior to experimental evolution, we washed bacterial pre-cultures twice with sterile NaCl (0.85%), adjusted OD_600_ to 1.0, and diluted 10^-4^ into 200 μL of iron-limited CAA (Bipyridyl 400 μM) per well in a 96-well plate. The plate was then incubated in a Tecan Infinite M-200 plate reader (Tecan Group Ltd., Switzerland) for 24 hours, and OD_600_ and pyoverdine-specific fluorescence (emission 400 nm, excitation 460 nm) were measured every 15 minutes.

### Experimental evolution

We conducted experimental evolution with *pvdS*_gene and *pvdS*_prom as starting points. We let each strain evolve independently under six different experimental treatments in a full-factorial design: 2 spatial structures (unstructured vs. structured) x 3 iron availabilities (low vs. medium vs. high iron availability) in three replicate independent lines (Fig. 2). At the start of the experimental evolution, overnight cultures of both clones were washed twice with NaCl (0.85%), adjusted to an OD600 of 1.0 and diluted 1:1000 into 200 μL of nutrient medium in 96-well plates. Plates were wrapped with parafilm, incubated for 24 hours and subsequently diluted 1:1000 in fresh nutrient medium. We repeated this cycle for 20 consecutive transfers, allowing for approximately 200 generations of bacterial evolution (Fig. 2). At the end of the experiment, we prepared freezer stocks for each evolved population (n = 36) by mixing 100 μL of bacterial culture with 100 μL of sterile glycerol (85%). Samples were stored at −80° C.

### Isolation of single clones

To check whether evolved clones showed altered pyoverdine production levels compared to the ancestral *pvdS*_gene and *pvdS*_prom strains, we isolated a total of 720 evolved clones (20 per replicate and treatment). Specifically, we regrew evolved bacterial populations from freezer stocks in 5 mL LB medium for 16-18 hours (180 rpm) and subsequently adjusted them to OD_600_ = 1.0. Then, 200 μL of 10^-6^ and 10^-7^ dilutions were spread on large LB agar plates (diameter 150 mm), which we incubated at 37° C for 18-20 hours. We then randomly picked twenty colonies for each of the 36 evolved populations, and immediately processed the clones for the pyoverdine measurement assay (see below).

### Screen for evolved pyoverdine production levels

For each of the 720 evolved clones, we transferred a small amount of material from the agar plate directly into 200 μL of CAA + Bipyridyl (400 μM) in individual wells on a 96-well plate. We incubated plates with clones originating either from unstructured environments or structured environments for 24 hours under shaken (180 rpm) or static conditions, respectively. Following incubation, we measured OD_600_ and pyoverdine-specific fluorescence (emission 400 nm, excitation 460 nm) in the Tecan Infinite M-200 plate reader as a single endpoint measurement. As controls, we included in three-fold replication on each plate: the high-producing PAO1 wildtype (positive control); the pyoverdine knockout mutant PAO1-Δ*pvdD*-*gfp* (negative control); the two low-producing mutants *pvdS*_gene and *pvdS*_prom; and blank growth medium. To preserve all tested clones for future experiments, we mixed 100 μL of bacterial culture with 100 μL of sterile glycerol (85%) for storage at −80° C.

### Confirmation of evolved pyoverdine phenotypes

Based on the screen above, we identified 34 clones with an altered pyoverdine production level (Table S1 in Additional File 1). Specifically, we found five clones that seem to have restored pyoverdine production by roughly 50% (i.e. in terms of the difference between the low-producing ancestor cheat and the high-producing wildtype) and 29 clones that seem to produce less than 33% of pyoverdine compared to their ancestral pyoverdine low-producers (either *pvdS*_gene or *pvdS*_prom). We subjected these clones to an in-depth repeated screening of their pyoverdine phenotype. In addition, we selected two random clones per treatment (n = 24), from different evolved populations, that displayed no change in their production levels (compared to *pvdS*_gene or *pvdS*_prom). One clone had to be excluded due to contamination, so that the final sample size for this group of clones was n = 23. For all of these evolved clones (n = 57), we re-measured their pyoverdine production level in three-fold replication using the same protocol and controls as described above.

### Sequencing of *pvdS* promoter and coding region

Since the ancestral low-producing strains (*pvdS*_gene or *pvdS*_prom) had mutations in the *pvdS* gene or its promoter, we were wondering whether the altered phenotypes observed in the evolved clones were based on reversion or additional mutations in this genetic region. To address this question, we PCR amplified and sequenced the *pvdS* gene and the upstream region containing the promoter sequence of all 57 evolved clones screened above. PCR mixtures consisted of 2 μl of a 10 μM solution of each primer, *pvdS_fw* (5’-GACGCATGACTGCAACATT-3’) and *pvdS_rev* (5’-CCTTCGATTTTCGCCACA-3’), 25 μl Quick-Load Taq 2X Master Mix (New England Biolabs), 1 μl of DMSO, and 20 μl of sterile Milli-Q water. We added bacterial biomass from glycerol stocks to the PCR mixture distributed in 96-well PCR plates. Plates were sealed with an adhesive film. We used the following PCR conditions: denaturation at 95° C for 10 min; 30 cycles of amplification (1 min denaturation at 95°C, 1 min primer annealing at 56°C, and 1 min primer extension at 72°C); final elongation at 72 °C for 5 min. The PCR products were purified and commercially sequenced using the *pvdS_fw* primer. While sequencing worked well for 51 clones, it failed for two clones, and resulted in partial sequences for six clones (Table S1 in Additional File 1).

### Statistical Analysis

All statistical analyses were performed using R 3.2.2[29]. We tested for treatment differences in the frequency of non-or low-producing strains using Fisher’s exact test and corrected for multiple testing using the Bonferroni correction. To compare pyoverdine production of evolved clones to that of the low-producing ancestors, we used one-way analyses of variance (ANOVA) and corrected for multiple testing using Tukey’s HSD (honest significant difference) test.

## Authors’ contributions

EG and RK planned the experiments. EG carried out the experiments and conducted statistical analysis. EG and RK analyzed and interpreted the data, and wrote the manuscript.

---

**Fig. S1:**
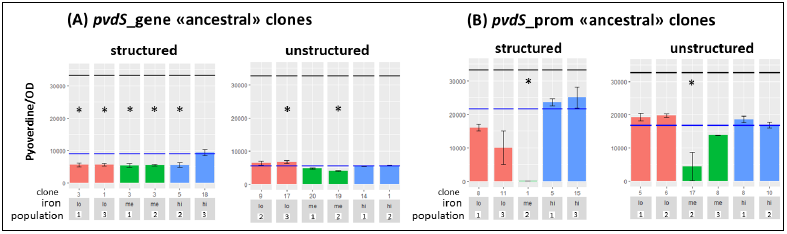
Confirmed pyoverdine phenotypes in selected clones. Evolved clones with ancestral pyoverdine production levels were re-tested to confirm their phenotype. Pyoverdine production was measured in iron-limited media. **(A)** Clones evolved from the low-producer *pvdS*_gene. **(B)** Clones evolved from the low-producer *pvdS*_prom. Y axes show pyoverdine-specific fluorescence divided by growth (optical density at 600 nm) after 24 h of incubation. X axes show independent replicate populations the clones evolved in and iron availability during experimental evolution. Bars represent mean values of three replicates per evolved clone. Error bars denote standard error of the mean. The black line represents the average wildtype production level in the same assay, while the blue line denotes the average production level of the respective low-producing ancestor. Bars are coloured by iron availability during evolution: red = low iron; green = medium iron; blue= high iron. We used one-way ANOVAs with Tukey’s post-hoc test for comparisons relative to the low-producing ancestor. Asterisks indicate a significant difference (p<0.05) from the ancestor.

**Table S1:**
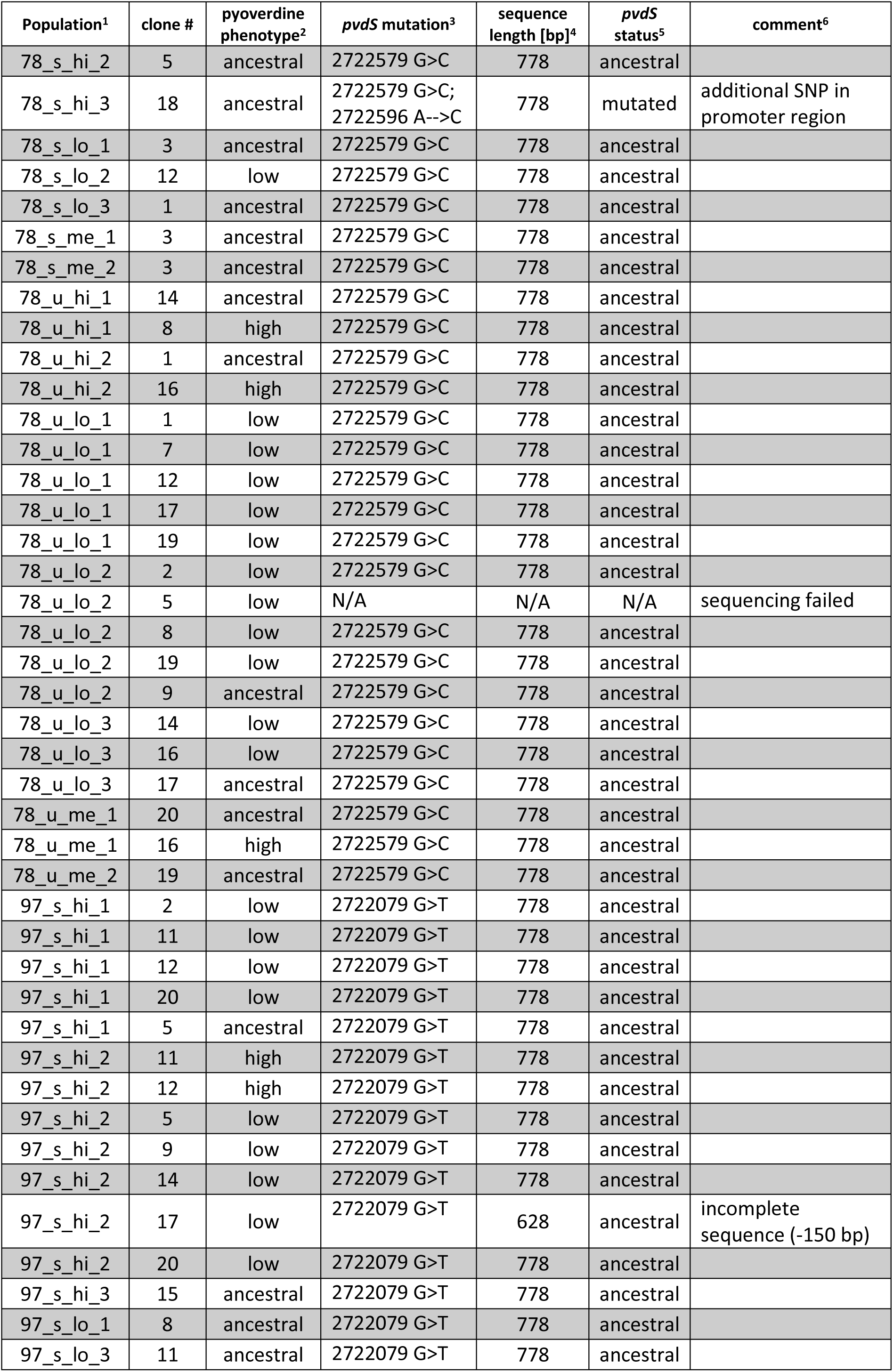
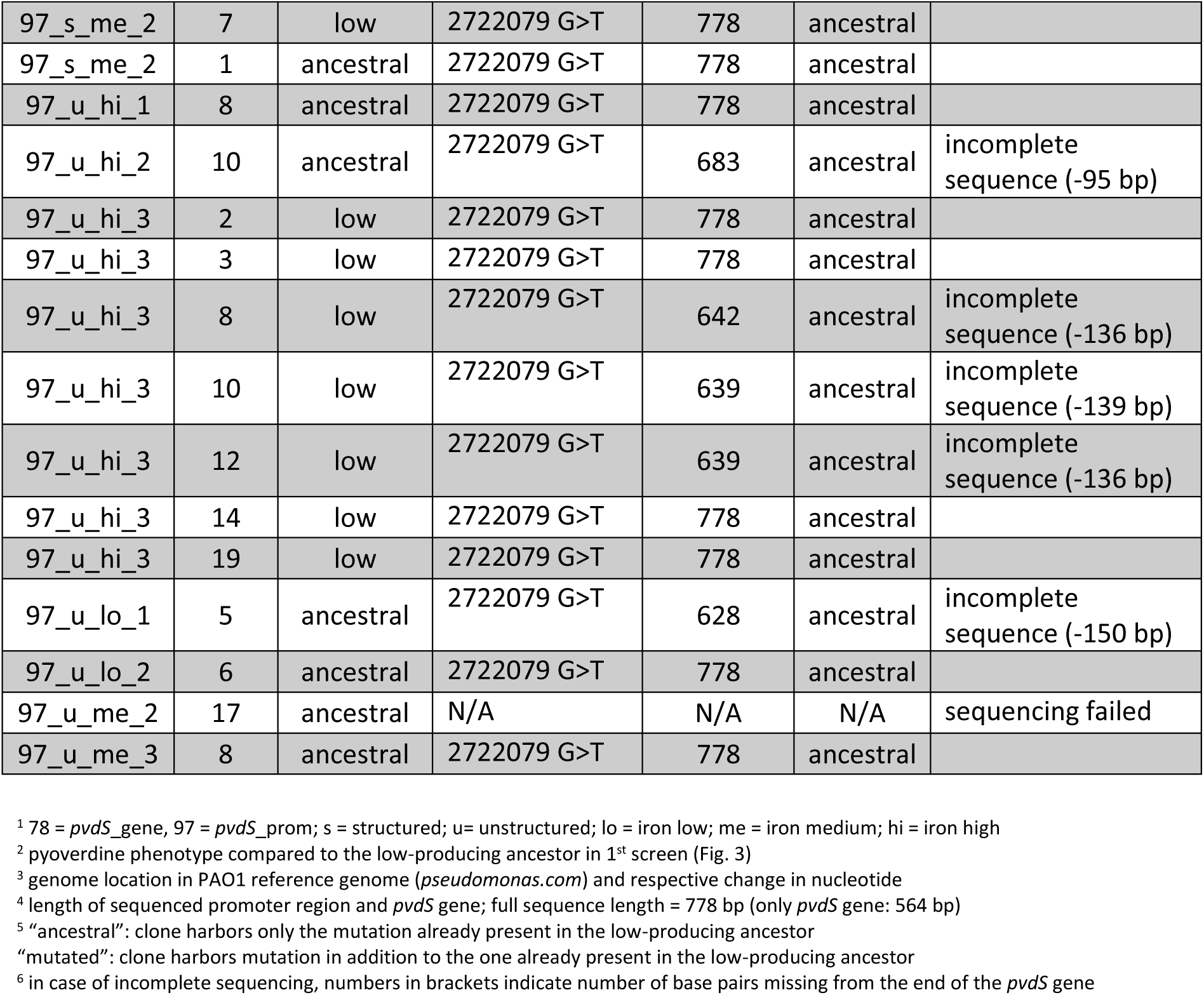
Clones selected for in-depth analysis and sequencing

